## Supplementary material for "Episodic long-term memory formation during slow-wave sleep": all supplements

Supplement

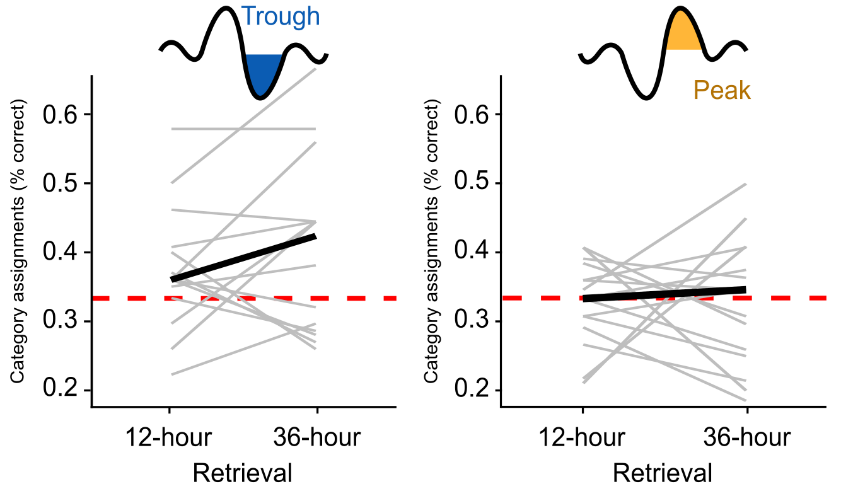

**Figure S1. Paired retrieval performance.**
Lines connecting participants’ retrieval performance after 12 hours and 36 hours (percentage correct category assignments). Results are faceted for the Trough (left, N=15) and the Peak (right, N=15) conditions. Thick black line depicts the group averages. The red dotted line represents chance-level performance. This is the same data as presented in Figure 1.

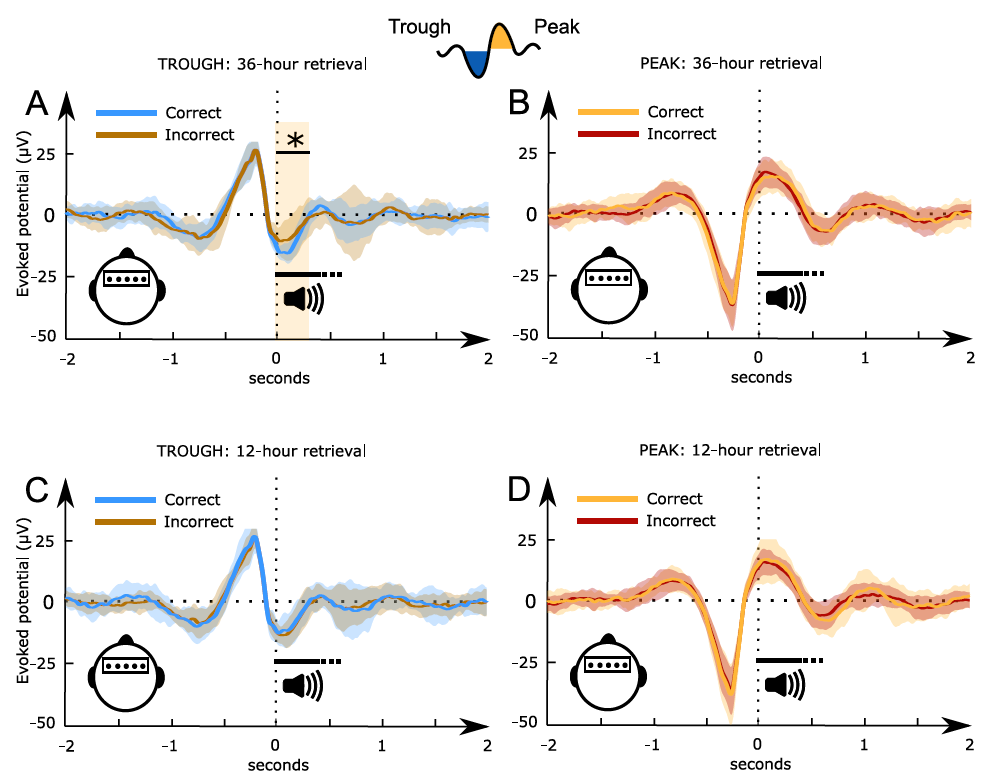

**Figure S2. Accuracy-related evoked EEG potentials recorded during sleep.** Comparison of word-related potentials measured during slow-wave sleep for later (12-hour retrieval in the bottom row and 36-hour retrieval in the top row) correctly and incorrectly assigned foreign words to superordinate categories. The left side displays voltage changes in relation to the targeted troughs (A, C) and the right side in relation to the target peaks (B, D). *, significant time point, cluster *p*< 0.05. Shaded areas represent the standard deviation between subjects. All trials are sorted with reference to the word onset (0 seconds, vertical dotted line).

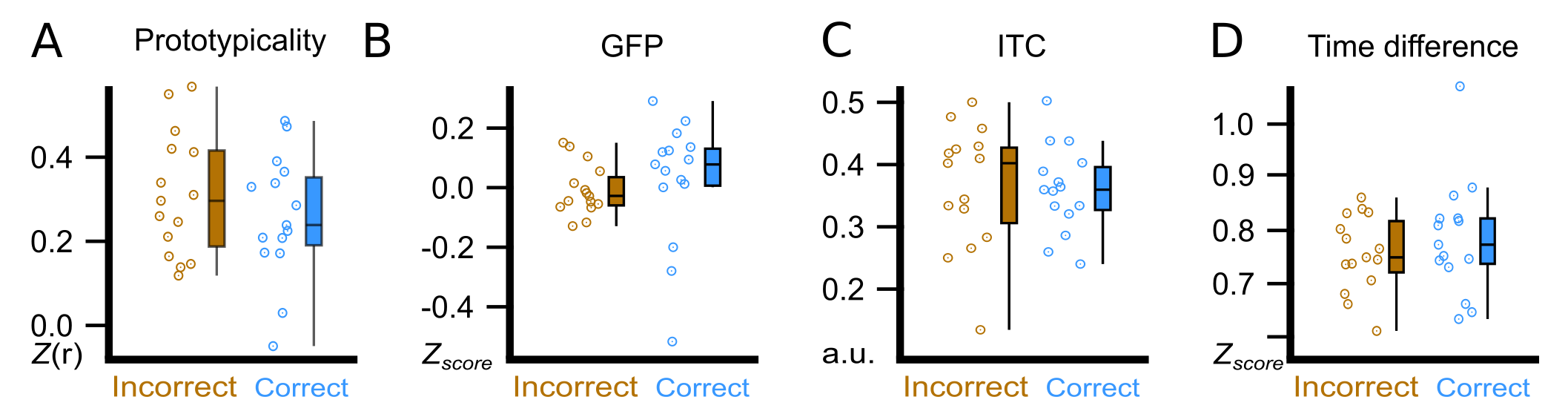

**Figure S3. Trough features compared between correct and incorrect category assignments at the 12-hour retrieval**. The same analysis as shown in Figure 4 of the manuscript, but for accuracy measured at the 12-hour retrieval. We display the comparison between correctly (blue) and incorrectly (brown) assigned foreign words in the experimental condition at the 12-hour retrieval using the amplitude of the correlation between the ERP and the template (Prototypicality) at word onset (**A**), using global field power (GFP) of the ERP at word onset (**B**), using the inter-trial phase coherence (ITC) between the four presentations of a word pair (**C**), and using the time difference (Time Difference) between the actual acoustic stimulation and the measured trough maximum (**D**). All paired t-test did not reveal a significant difference (p>0.15). Participant averages (dots), group averages (vertical black line) and 95% confidence intervals (boxes) are displayed.

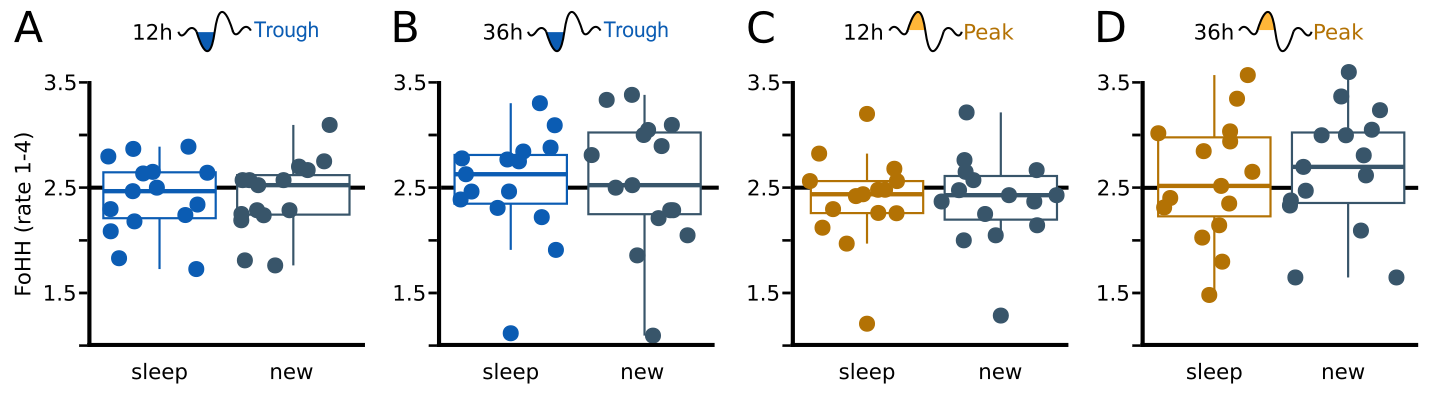

**Figure S4. Feeling of having heard (FoHH) for sleep-paired and new pseudowords**. Boxplot of subject averages of FoHH, faceted for sleep-paired (blue or brown) and new (grey) pseudowords. The conditions are from left to right: **(A)** Trough condition, 12h retrieval (blue); **(B)** Trough condition, 36h retrieval (blue); **(C)** Peak condition, 12h retrieval (brown); **(D)** Peak condition, 36h retrieval (brown). The FoHH did not significantly differ between sleep-heard and new pseudowords in all four conditions (all p>0.1). Similarly, an ANOVA of the sleep-paired pseudowords did not yield any significant main effect of stimulation (Peak & Trough), time of retrieval (12h & 24h) or the interaction (all p<0.1). The dots represent each subject’s average.

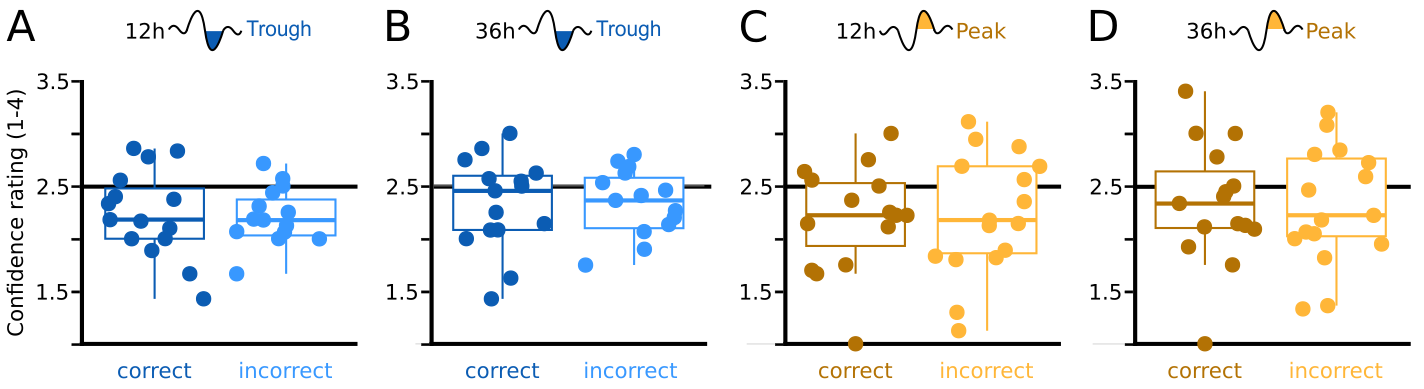

**Figure S5. Confidence ratings of category assignments, separated by retrieval accuracy**. Confidence ratings did not differ between correctly or incorrectly categorized pseudowords (all p<0.5): **(A)** in the Trough condition after 12h (blue, left), **(B)** in the Trough condition after 36h (blue, right), **(C)** in the Peak condition after 12h (brown, left) and **(D)** in the Peak condition after 36h (brown, right). Confidence ratings ranged from one to four. An ANOVA, also including the ratings of new pseudowords, did not yield a main effect of stimulation (Peak & Trough), time of retrieval (12h & 24h), presentation (sleep-heard, new) or any interaction (all p<0.2).

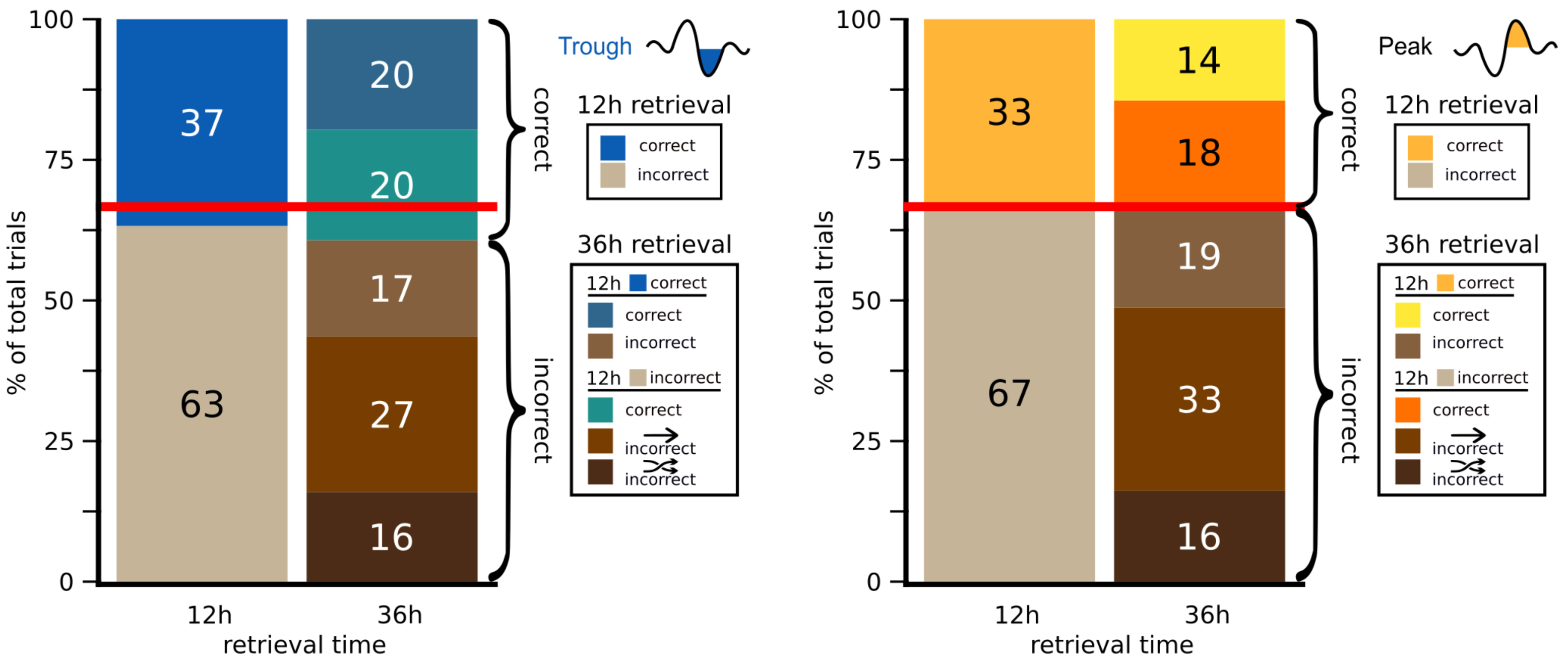

**Figure S6. Switching rate of category assignment.** On the left, we present the trial-based answer consistency for the Trough condition. Percentage of trials that were assigned to the correct (blue-ish) and incorrect (brown-ish) superordinate category, based on the sleep learned association. Category assignment accuracy is depicted for the retrieval after 12h (left) and for the retrieval after 36h (right). The correct responses after 36h are either after a correct (dark blue) or an incorrect (turquoise) assignment at 12 hours. Incorrect responses either came from a previously correct assignment (light brown), the same incorrect category assignment (brown) or the other incorrect category (dark brown). Similarly, on the right, we present the switching rate for the Peak condition. Orange-ish are here correctly assigned trials. Please note that at the 36h retrieval only 14% were twice correct (yellow, top square), compared to 20% in the Trough condition. Red line represents chance performance to choose an incorrect category that is two out of three categories (0.66%). Straight arrow = consistent incorrect category assignment (e.g. response twice ‘animal’, correct = tool), crossed arrows = inconsistent incorrect category assignment (e.g. correct = building, 12h response = animal, 36h response = tool).
